# Loss of ovarian function and estrogen therapy remodel the brain’s synaptic and metabolic proteome

**DOI:** 10.1101/2025.10.20.683488

**Authors:** Sebastian F. Salathe, Edziu Franczak, Zane Busick, Frederick B. Boakye, Julie Allen, Andrew Lutkewitte, Heather Wilkins, Thyfault John, Benjamin A. Kugler

**Affiliations:** Department of Cell Biology and Physiology, University of Kansas Medical Center, Kansas City KS, USA; Internal Medicine, Division of Endocrinology and Clinical Pharmacology and KU Diabetes Institute, University of Kansas Medical Center, Kansas City, KS, USA; Kansas Center for Metabolism and Obesity Research, University of Kansas Medical Center, Kansas City, KS, USA; The University of Kansas Diabetes Institute, Kansas City KS, USA; Department of Neurology, University of Kansas Medical Center, Kansas City KS, USA; University of Kansas Alzheimer’s Disease Center, University of Kansas Medical Center, Kansas City KS, USA; Department of Biochemistry and Molecular Biology, University of Kansas Medical Center, Kansas City KS, USA

## Abstract

Menopause is linked to cognitive decline and reduced brain metabolism, while estrogen (E2) therapy has been shown to mitigate these effects. Understanding the molecular mechanisms by which ovarian hormones and E2 influence neuroprotection is essential for developing strategies to maintain brain health in women. In this study, we examined how the loss of ovarian hormones, with or without E2 treatment, affects the brain proteome and mitochondrial energy production in aged female C57BL/6J mice (36–40 weeks). The mice underwent sham or ovariectomy (OVX) surgery and were fed a high-fat diet for 10 weeks; six weeks after surgery, OVX mice received either sesame oil or E2 treatment for four weeks. Proteomic analysis of brain homogenates revealed 4,992 proteins regulated by E2, with pathway analysis showing increased signaling proteins related to synaptogenesis. OVX reduced proteins involved in synaptic function, branched-chain amino acid and ketone metabolism, the TCA cycle, and oxidative phosphorylation (Complexes I, IV, and V), while E2 restored protein expression within these pathways. Despite alterations in OxPhos proteins, basal and state 3 mitochondrial respiration remained unchanged, although notable impairments to Complex IV enzymatic activity were apparent in OVX, but not following E2 replacement. Overall, these results indicate that E2 supports brain health by maintaining proteins crucial for synaptic integrity and metabolism, and by reducing the decline in mitochondrial bioenergetics associated with menopause.

## INTRODUCTION

Women experience a higher incidence of cognitive decline with aging and are more predisposed to neurological diseases such as Alzheimer’s disease (AD), a disparity increasingly linked to the hormonal transition of menopause. Compared to age-matched men, peri – and post-menopausal women exhibit accelerated cognitive decline, greater brain atrophy, and reduced brain glucose metabolism [1, 2]. Preclinical ovariectomy (OVX) studies, which model the abrupt loss of ovarian hormones, similarly demonstrate impairments in memory, synaptic signaling, and mitochondrial function [3, 4]. Together, these findings suggest that estrogen loss contributes to brain vulnerability during aging by disrupting neuroplasticity and metabolic processes that support cognitive health.

Hormone therapy (HT), via pharmacological treatment with exogenous estrogen and progesterone, has been associated with slower cognitive decline, reduced Aβ burden, and preserved brain volume [5–13]. Estrogen (E2) specifically has been shown to enhance neuroplasticity by upregulating growth-associated proteins [14], preserve cholinergic function through increased choline acetyltransferase (ChAT) activity and acetylcholine release [12, 15], and support bioenergetics by maintaining brain glucose metabolism and mitochondrial respiration [16, 17]. At the molecular level, E2 influences the brain proteome, increasing synaptic proteins such as syntaxin binding protein 1 (Stxbp1), Snap25, and dynamin [18], while also modulating mitochondrial proteins within the tricarboxylic acid (TCA) cycle and ATP synthase [16]. However, prior proteomic studies using 2D-DIGE lacked sufficient depth to capture the full scope of E2- mediated regulation of the brain proteome. Moreover, these studies often used rodents at ages not reflective of the menopausal transition while examining only acute (24 h) E2 treatment, thereby capturing rapid rather than long-term proteomic changes.

To address this gap, the present study employed whole-brain homogenates and an unbiased proteomic approach to define how OVX and E2 treatment remodel the brain proteome in an aged mouse model. We hypothesized that E2 would restore protein expression in pathways critical for metabolism, synaptic plasticity, and mitochondrial bioenergetics that would be downregulated in female OVX mice. Our data reveal that E2 replacement rescues proteins involved in metabolic pathways (glucose, branched-chain amino acids, ketones), mitochondrial bioenergetics (TCA cycle, Complex IV, Complex V), and synaptic signaling (Stxbp1, Snap25, Stx1a), all of which were reduced following OVX. These findings provide new mechanistic insight into how E2 supports brain health following the transition to menopausal and provide molecular targets for therapeutic intervention in cognitive decline.

## METHODS

### Animals

Female C57BL/6J mice carrying a floxed estrogen receptor alpha (ERα) allele, bred in-house at the University of Kansas Medical Center, were co-housed at thermoneutral temperature (30 °C) on a reverse 12:12 h light-dark cycle. Mice had ad libitum access to water and a low-fat diet (LFD; Research Diets D12450H, 10% kcal fat, 17% kcal sucrose) until initiation of experimental treatments. All procedures were approved by the Institutional Animal Care and Use Committee of the University of Kansas Medical Center.

### Experimental Treatment

At 36–40 weeks of age, female C57BL/6J mice underwent either sham surgery or bilateral ovariectomy (OVX) to deplete ovarian hormone production (n=9-11 per group). Immediately following surgery, all the mice were transitioned to a high-fat diet (HFD; Research Diets D12451, 45% kcal fat, 17% kcal sucrose, 1% cholesterol wt./wt., 4.68 kcal/g) for 10 weeks. The HFD plus OVX combined treatment was designed to replicate worsened metabolic health that can be induced by menopause clinically [19, 20].

### Estrogen Replacement

After 6 weeks of HFD feeding, OVX mice underwent surgical implantation of silastic tubing (20 mm length, sealed with surgical glue) between the scapulae, filled with either sesame oil or 17β-estradiol (E2; 36 μg/mL sesame oil), as previously described [21, 22]. To prevent hormone depletion, silastic tubing was replaced after 2 weeks. Sham-operated mice received sesame oil implants following this same paradigm.

### Anthropometrics and Tissue collection

Body weight was recorded weekly throughout the study. Body composition was measured via magnetic resonance imaging (EchoMRI, Houston, TX) at baseline after surgery or sham (prior to E2 replacement) and the day of euthanasia. Fat-free mass was calculated as total body mass minus fat mass.

At study termination, mice were anesthetized with phenobarbital (0.5 mg/g body weight) following a 2-hour fast (7:00–9:00 am). Blood was collected via cardiac puncture, allowed to clot at room temperature, placed on ice for 10 minutes, and centrifuged at 7,000 × g for 10 minutes at 4 °C to isolate serum. Brains were rapidly dissected: the right hemisphere was flash-frozen in liquid nitrogen for later processing, while the left hemisphere was used for mitochondrial isolation. Uterine wet weights were recorded as a marker of estrogen status.

### Mitochondrial Isolation

Following excision, the right hemisphere was placed in 8 mL of ice-cold mitochondrial isolation buffer (220 mM mannitol, 70 mM sucrose, 10 mM Tris, 1 mM EDTA, pH at 7.4) and homogenized using a Teflon pestle. Dounce glass-on-glass homogenization in combination with density centrifugation was performed as previously described to generate a mitochondrial-enriched fraction [23]. Briefly, brain homogenate was centrifuged at 1,500 x g (10 min at 4°C) and the supernatant filtered through cheese cloth before being pelleted (8,000 x g, 10min at 4°C). The pellet was then resuspended in 6 mL of mitochondrial isolation buffer, before being pelleted again at 6,000 x g (10min at 4°C). Following resuspension in 4 mL of mitochondrial isolation buffer containing 0.1% fatty acid-free BSA, mitochondrial fractions were spun at 4,000 x g (10min at 4°C). The final isolated mitochondrial pellet was resuspended in 350-400 μL of modified MiR05 mitochondrial respiration buffer (0.5 mM EGTA, 3 mM MgCl2, 60 mM KMES, 20 mM glucose, 10 mM KH2PO4, 20 mM HEPES, 110 mM sucrose, and 0.1% fatty acid-free BSA, pH adjusted to 7.1). Bicinchoninic acid assay was performed to determine the protein concentration of the isolated mitochondrial fraction.

### Mitochondrial Respiration

Real-time mitochondrial oxygen consumption (JO₂; pmol·s⁻¹·mL⁻¹) was measured using the Oroboros O2k high-resolution respirometer (Oroboros Instruments, Innsbruck, Austria) as previously described [24]. Following air calibration, isolated mitochondria were added to chambers containing 2 mL of modified MiR05 respiration buffer (37 °C) supplemented with 2 mM malate, 63.5 μM free CoA, and 2.5 mM L-carnitine, and continuously stirred at 750 rpm. Glutamate-supported respiration was initiated with 2 mM glutamate, followed sequentially by the addition of 2.5 mM ADP (State 3), 10 mM succinate (state 3S), 2 μM rotenone (complex I inhibitor), 5 mM malonate (complex II inhibitor), and 2 mM ascorbate + 0.5 mM TMPD (complex IV substrate). Complex I activity was calculated as State 3S - rotenone, and complex II activity as rotenone - malonate. Data were acquired and analyzed with DatLab software (version 7, Oroboros Instruments) and normalized to total mitochondrial protein content in each chamber.

### Proteomics

Brain proteomics were performed as previously described [25]. In brief, brain lysates were employed for proteomics analysis, following established protocols [26, 27]. Proteins underwent digestion using sequencing-grade modified porcine trypsin (Promega). Subsequently, peptides were separated on an in-line 150 x 0.075 mm column packed with reverse phase XSelect CSH C18 2.5μm resin (Waters), utilizing an UltiMate 3000 RSLCnanosystem (Thermo). Elution was achieved through a 60-minute gradient from a 98:2 to 65:35 buffer A:B ratio (buffer A = 0.1% formic acid, 0.5% acetonitrile; buffer B = 0.1% formic acid, 99.9% acetonitrile). The eluted peptides were ionized by electrospray (2.2kV) and subjected to mass spectrometric analysis on an Orbitrap Exploris 480 mass spectrometer (Thermo).

To generate a chromatogram library, six gas-phase fractions were acquired on the Orbitrap Exploris, with 4m/z DIA spectra utilizing a staggered window pattern from narrow mass ranges and optimized window placements. Precursor spectra were obtained after each DIA duty cycle, covering the m/z range of the gas-phase fraction. For wide-window acquisitions, the Orbitrap Exploris was configured to acquire a precursor scan followed by 50 times 12m/z DIA spectra using a staggered window pattern with optimized placements. Precursor spectra were again acquired after each DIA duty cycle.

Proteomic data underwent search using an empirically corrected library, followed by a quantitative analysis to obtain a comprehensive proteomic profile. Identification and quantification were performed using EncyclopeDIA and visualized with Scaffold DIA, incorporating 1% false discovery thresholds at both the protein and peptide levels. The obtained proteomics data were cross-referenced with the MitoCarta3.0 gene inventory to pinpoint proteins with robust evidence of mitochondria localization [28]. Any proteins not included within the MitoCarta3.0 gene inventory were excluded from subsequent analyses. Qiagen’s Ingenuity Pathway Analysis (IPA) (41) was executed to discern changes in global and mitochondrial pathways. Employing expression core analysis, with the expression log ratio (logfold change) incorporated for each respective protein, the directionality (Z-score) of pathway regulation was determined.

### Statistics

Data are presented as mean ± SD. Group differences were assessed using one-way ANOVA, followed by Tukey’s post hoc test when appropriate. Outliers were identified and removed using Grubb’s test. All analyses were conducted in GraphPad Prism 10, with statistical significance set at *P* < 0.05.

## RESULTS

### Estrogen treatment reduces body mass and fat mass in OVX mice

Six weeks following OVX and HFD, there were no differences in body, fat (FM), and fat-free mass (FFM) between the sham, OVX, and OVX+E2 groups (**Table 1**). However, OVX tended to increase fat and body mass compared to the sham group following 10 weeks of a HFD. Four weeks of E2 treatment (OVX+E2) reduced body mass and FM compared to the untreated OVX counterpart (*P*<0.05). Confirming OVX and E2 treatment, the uterus mass was significantly reduced in the OVX group; however, E2 treatment increased the uterus mass compared to both the Sham and OVX groups (*P*<0.05).

**Table 1.**
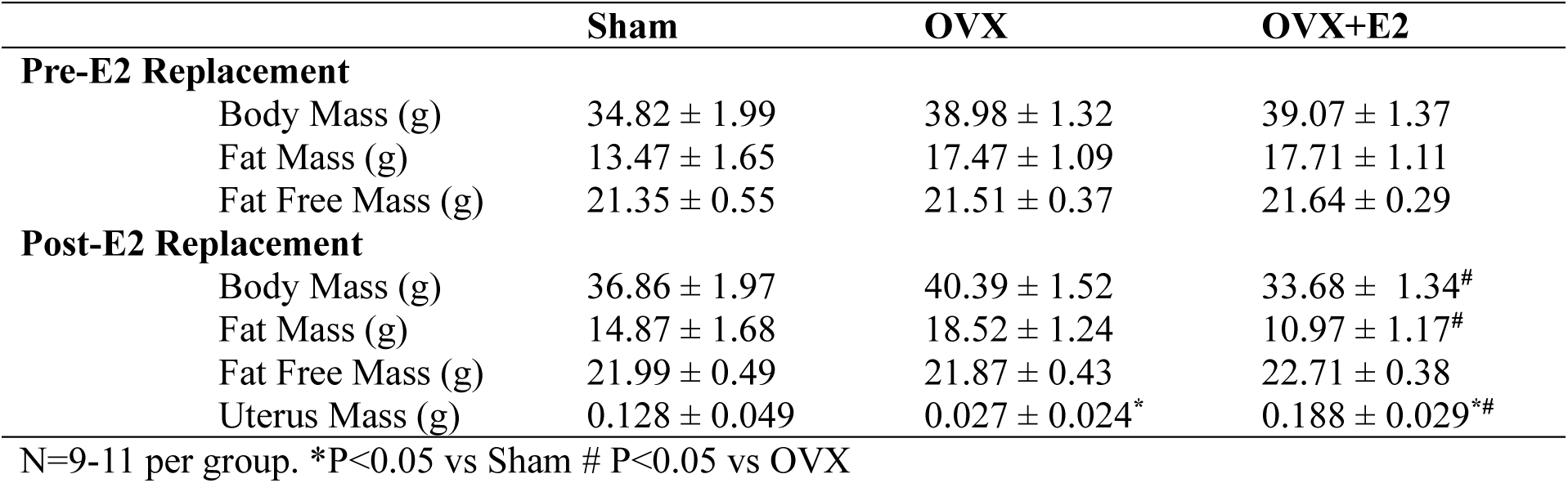
Anthropometric.

|  | **Sham** | **OVX** | **OVX+E2** |
| --- | --- | --- | --- |
| **Pre-E2 Replacement** |  |  |  |
| Body Mass (g) | 34.82 ± 1.99 | 38.98 ± 1.32 | 39.07 ± 1.37 |
| Fat Mass (g) | 13.47 ± 1.65 | 17.47 ± 1.09 | 17.71 ± 1.11 |
| Fat Free Mass (g) | 21.35 ± 0.55 | 21.51 ± 0.37 | 21.64 ± 0.29 |
| **Post-E2 Replacement** |  |  |  |
| Body Mass (g) | 36.86 ± 1.97 | 40.39 ± 1.52 | 33.68 ± 1.34<sup>#</sup> |
| Fat Mass (g) | 14.87 ± 1.68 | 18.52 ± 1.24 | 10.97 ± 1.17<sup>#</sup> |
| Fat Free Mass (g) | 21.99 ± 0.49 | 21.87 ± 0.43 | 22.71 ± 0.38 |
| Uterus Mass (g) | 0.128 ± 0.049 | 0.027 ± 0.024<sup>*</sup> | 0.188 ± 0.029<sup>*#</sup> |

N=9-11 per group. \*P<0.05 vs Sham # P<0.05 vs OVX

### Estrogen treatment enriches the proteins associated with synaptogenesis signaling

Proteomic profiling identified 7,781 unique proteins across all groups, with no overall difference in total abundance between groups (**Figure 1A**). E2-regulated proteins were defined as those altered by OVX (OVX vs. sham) and reversed by E2 treatment (OVX+E2 vs. OVX), yielding 4,992 proteins (**Figure 1B**). Pathway analysis reveals OVX increased proteins involved in oxidative phosphorylation, mitochondrial protein degradation, and ribosomal quality control (**Figure 1C**), whereas E2 treatment in the OVX condition increased pathways involved in synaptogenesis, glutamatergic receptor signaling, and protein ubiquitination (**Figure 1D**). Several pathways were reciprocally altered between OVX and E2 treatment groups, suggesting that E2 replacement reverses many of the proteomic changes induced by estrogen loss. Specifically, pathways such as synaptogenesis signaling, GABA receptor signaling, and LXR/RXR activation, which were suppressed or dysregulated by OVX, were restored toward sham levels following E2 treatment (**Figure 1E**). This indicates that chronic E2 treatment reestablishes a protein expression profile resembling that of intact, estrogen-sufficient mice.

**Figure 1:**
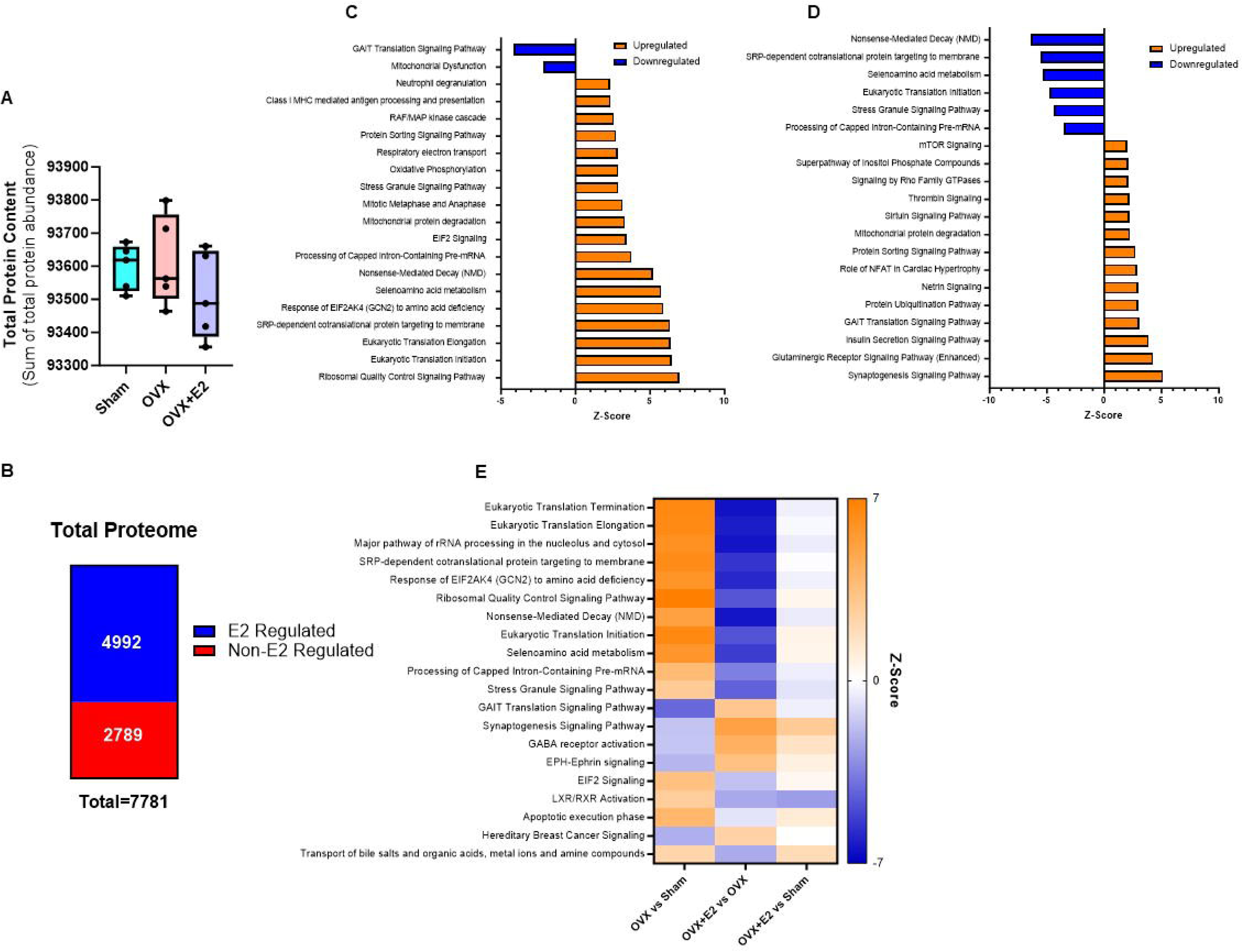
Whole-brain proteomic analysis and pathway enrichment. (**A**) Total proteome abundance. (**B**) Proteins divergently regulated by E2 treatment. (**C**) Top 20 pathways altered in OVX vs. Sham mice identified by IPA. (**D**) Top 20 pathways altered in OVX+E2 vs. OVX identified by IPA. (**E**) Top 20 divergent pathways regulated by E2 treatment. Data are presented as mean ± SD. *n* = 5 mice per group.

### Estrogen treatment after OVX restores presynaptic vesicle trafficking and synaptic signaling molecule proteins

Focusing on the synaptogenesis signaling pathway, we identified 249 out of 316 proteins, of which 158 were specifically regulated by E2 treatment after OVX (**Figure 2A**). The overall summed abundance of these divergent proteins did not differ between groups (**Figure 2B**). We found that synaptogenesis proteins upregulated by E2 were significantly reduced in OVX mice and restored by E2 treatment **(Figure 2C**, *P*<0.05). In contrast, the total abundance of proteins downregulated by E2 showed no group differences (**Figure 2D**). Because the downregulated proteins were unchanged, we focused subsequent analyses on the upregulated subset, which captured the significant effects of OVX and E2 treatment. We further classified these proteins into four functional groups: presynaptic vesicle trafficking, synaptic cellular signaling molecules, postsynaptic receptors/ion channels, and synaptic adhesion. OVX significantly decreased the total abundance of proteins in presynaptic vesicle trafficking and synaptic cellular signaling, both of which were restored with E2 treatment (**Figures 2E–F**, *P*<0.05). Similarly, in postsynaptic receptors/ion channels, the total protein abundance was increased by E2 treatment compared to OVX (**Figure 2G**, *P*<0.05), while no significant changes were observed in the total abundance of synaptic adhesion proteins (**Figure 2H**). Heatmap analyses highlighted individual proteins driving these changes, including Syt7, Rab3a, Rab5c, Snap25, and Stxbp1 within presynaptic vesicle trafficking (**Figure 2I**, *P*<0.05), Prkcd and Pik3ca among synaptic signaling molecules (**Figure 2J**, *P*<0.05), Grm8 in postsynaptic receptors (**Figure 2K**, *P*<0.05), and EphA6 within synaptic adhesion (**Figure 2L**, *P*<0.05).

**Figure 2:**
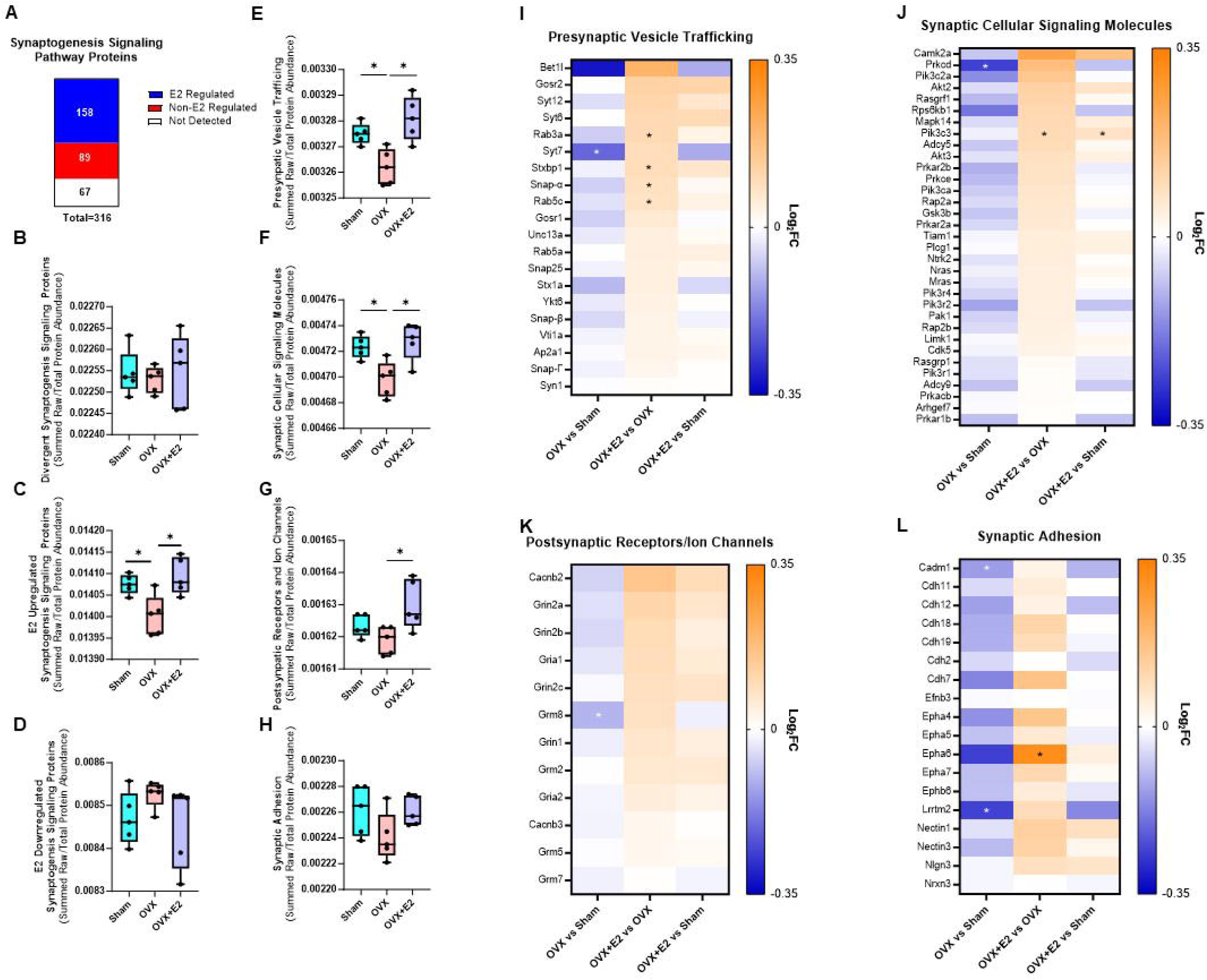
Synaptogenesis Pathway Analysis. (**A**) Synaptogenesis signaling proteins divergently regulated by E2. Protein abundance of E2-upregulated (**B**) and E2-downregulated (**C**) proteins. Total protein abundance of E2-upregulated synaptic categories: (**D**) presynaptic vesicle proteins, (**E**) synaptic cellular signaling molecules, (**F**) postsynaptic receptors and ion channels, and (**G**) synaptic adhesion proteins. (**H-K**) Heatmap of log₂ fold-change for individual proteins across synaptic categories. Data are presented as mean ± SD. *n* = 5 mice per group. * *P* < 0.05 vs. indicated group.

### Brain mitochondrial protein content is reduced with OVX and restored following estrogen treatment

To examine mitochondrial alterations driven by OVX and E2 treatment, we cross-referenced our proteomic dataset with MitoCarta3.0, identifying 887 out of 1140 mitochondrial-associated proteins (**Figure 3A**). Of the 887 mitochondrial-associated proteins, E2 treatment after OVX regulated 515 proteins (**Figure 3A**). Summed mitochondrial protein abundance was significantly reduced by OVX relative to sham and restored with E2 treatment (**Figure 3B**, *P*<0.05). Pathway enrichment of the MitoCarta3.0 subset revealed that OVX increased proteins involved in mitochondrial protein degradation, complex IV assembly, TCA cycle regulation, and estrogen receptor signaling (**Figure 3C**). In contrast, E2 treatment after OVX enriched proteins involved in glutathione redox reactions, mitochondrial translation, complex IV assembly, and peroxisomal lipid metabolism, while downregulating mitochondrial calcium ion transport (**Figure 3D**). Direct comparison of pathways altered by OVX and E2 treatment revealed opposing regulation of multiple mitochondrial functions, including TCA cycle regulation, complex IV assembly, branched-chain amino acid catabolism, and fatty acyl-CoA biosynthesis (**Figure 3E**).

**Figure 3:**
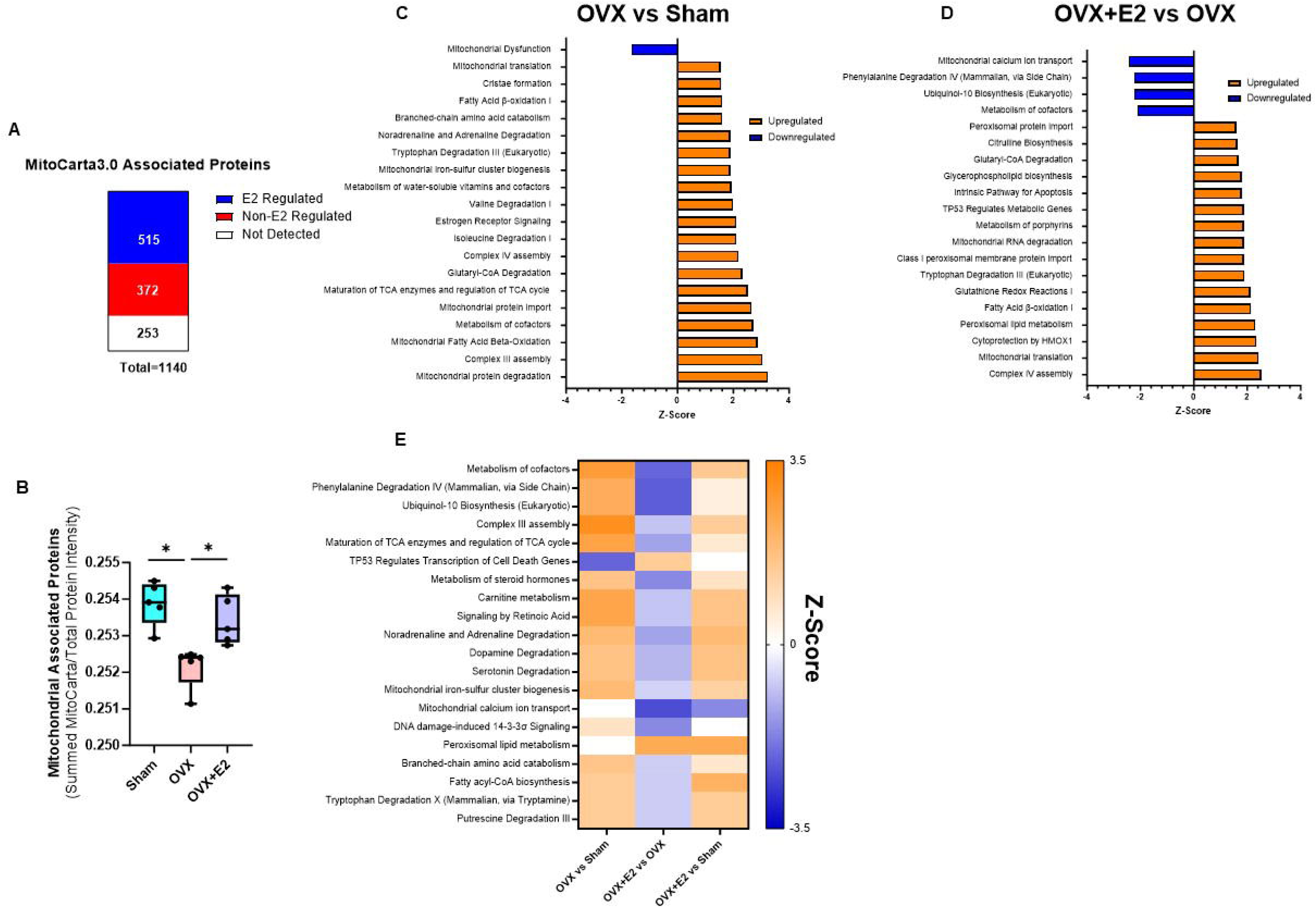
MitoCarta3.0 proteomic analysis and pathway enrichment. (**A**) MitoCarta3.0 proteins divergently regulated by E2 treatment. (**B**) MitoCarta3.0 total protein abundance. (**C**) MitoCarta3.0 top 20 pathways altered in OVX vs. Sham mice identified by IPA. (**D**) MitoCarta3.0 top 20 pathways altered in OVX+E2 vs. OVX identified by IPA. (**E**) MitoCarta3.0 Top 20 divergent pathways regulated by E2 treatment. Data are presented as mean ± SD. *n* = 5 mice per group.* *P* < 0.05 vs. indicated group.

### Estrogen treatment regulates proteins associated with metabolism and OXPHOS

To examine the effects of E2 on brain metabolism, we assessed proteins from key metabolic pathways, including glycolysis, fatty acid oxidation, branched-chain amino acid (BCAA) metabolism, and ketone metabolism. No significant differences were observed in glycolytic protein levels between groups **(Figure 4A**). In contrast, mitochondrial pyruvate transport and lactate dehydrogenase levels were significantly increased with E2 replacement after OVX compared to OVX alone (**Figures 4B-C**; *P*<0.05). There was no overall change in summed protein levels associated with β-oxidation across groups (**Figure 4D**). OVX significantly lowered the levels of proteins involved in BCAA metabolism and ketone metabolism, both of which were restored by E2 treatment (**Figures 4E-F**; *P*<0.05). Similarly, TCA cycle protein levels were reduced following OVX and recovered with E2 treatment after OVX (**Figure 4G**; *P*<0.05). To visualize these changes, heatmaps were created showing the log2 fold change (log2FC) of proteins involved in ketone metabolism, glycolysis, BCAA metabolism, β-oxidation, and the TCA cycle (**Figure 4H-L**).

**Figure 4:**
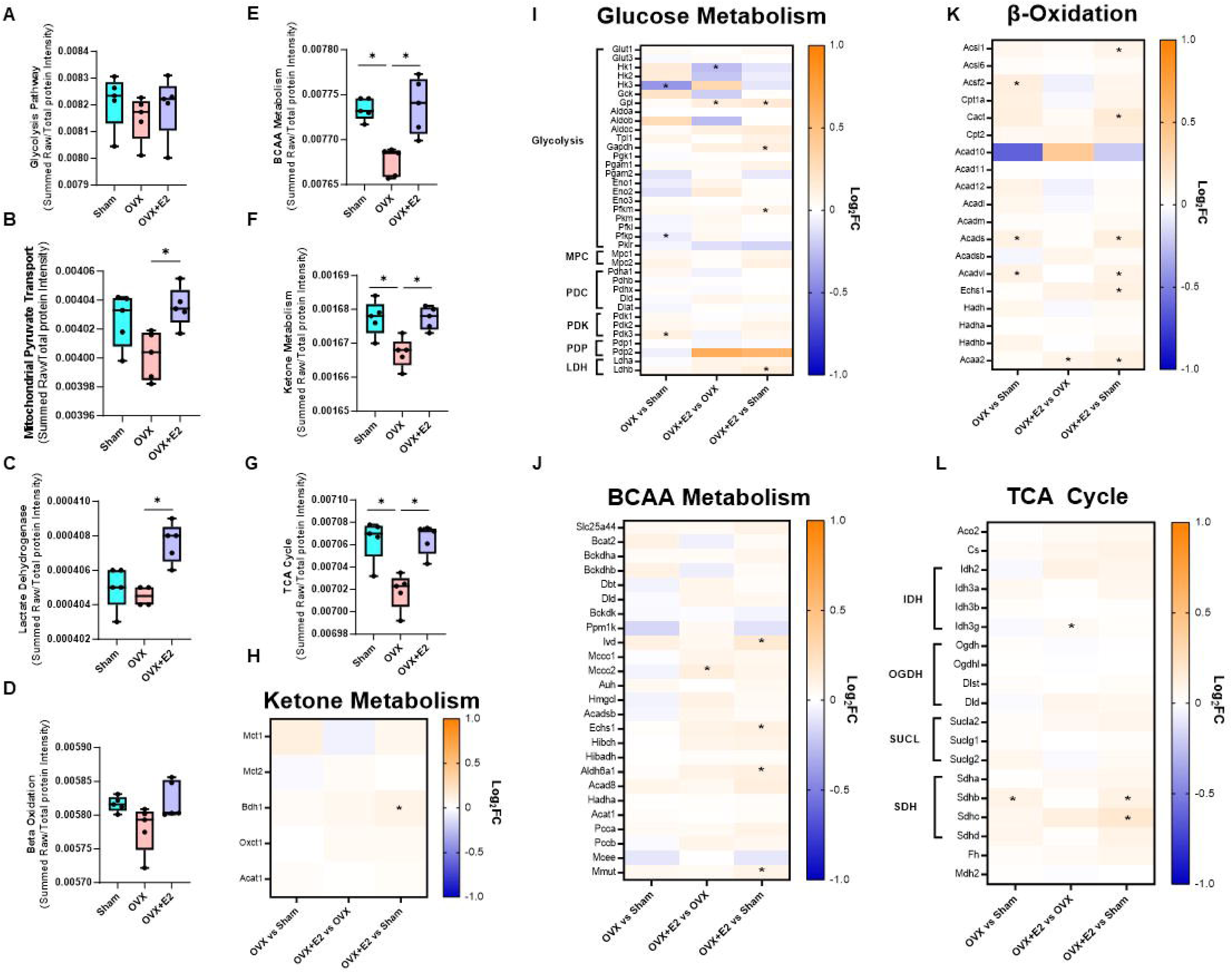
Metabolism pathway analysis: Total protein abundance of (**A**) glycolysis, (**B**) mitochondrial pyruvate transport, (**C**) lactate dehydrogenase, (**D**) fatty acid beta oxidation, (**E**) branch chain amino acid metabolism, (**F**) ketone metabolism, and (**G**) TCA cycle. (**H-L**) Heatmap of log₂ fold-change for individual proteins across metabolism categories. Data are presented as mean ± SD. *n* = 5 mice per group. * *P* < 0.05 vs. indicated group.

We next evaluated protein abundance across the oxidative phosphorylation (OXPHOS) complexes. OVX reduced the abundance of Complex I, IV, and V proteins (**Figure 5A, D, E**; *P*<0.05). E2 treatment selectively restored Complex V and increased Complex II protein abundance compared to OVX (**Figure 5B, E**; *P*<0.05), while Complex III proteins remained unchanged (**Figure 5C**). To examine subunit-specific effects, we generated a heatmap of log2FC (**Figure 5F**). Within Complex I, OVX decreased subunits Ndufb2 and Ndufs5, both of which were restored with E2 treatment (*P*<0.05). In Complex IV, E2 increased the abundance of subunits Cox15 and Cox20, whereas in Complex V, OVX reduced Atp5pf, while E2 elevated Atp5mj (*P*<0.05).

**Figure 5:**
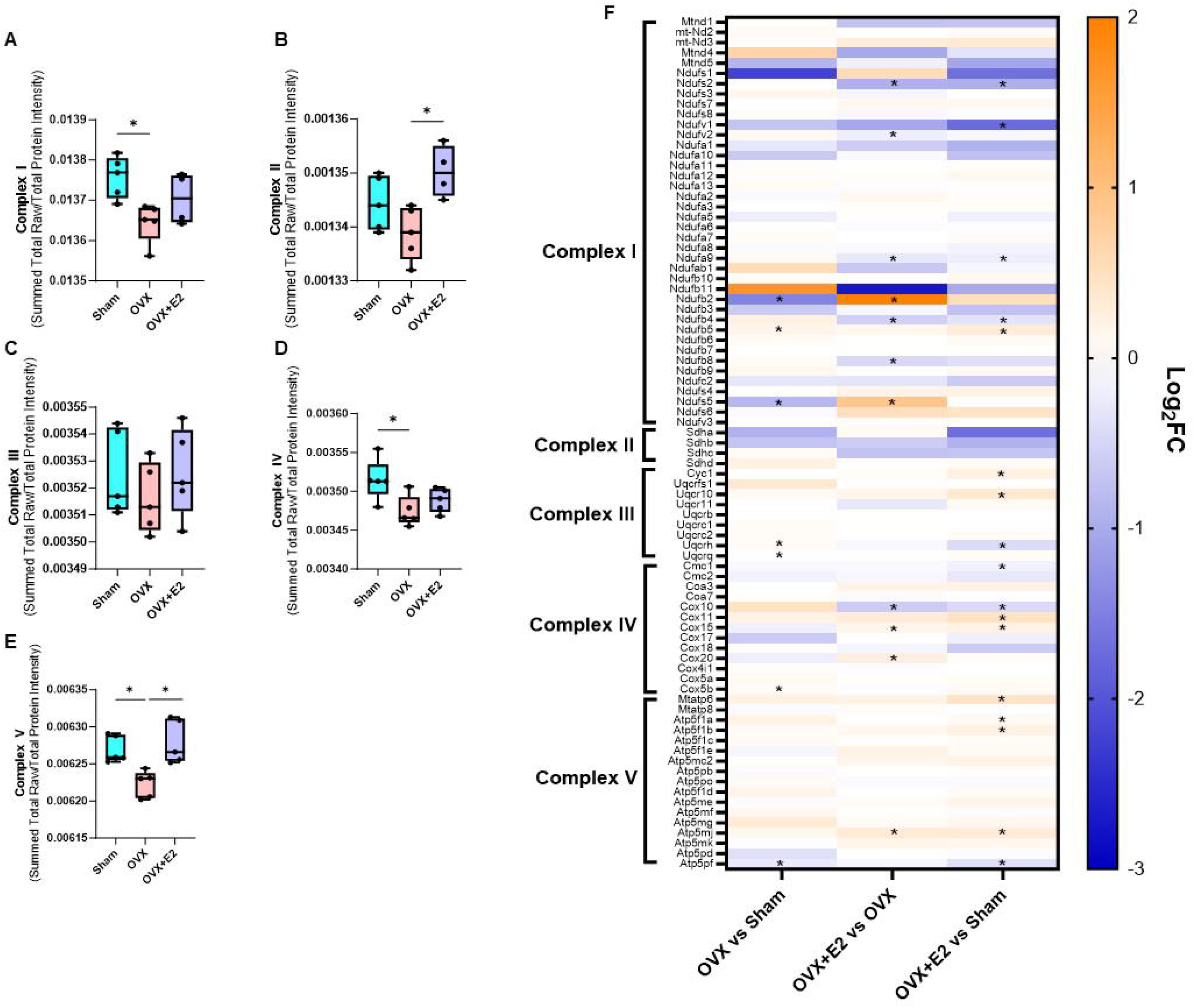
Oxidative phosphorylation protein analysis: Total protein abundance of (**A**) Complex I, (**B**) Complex II, (**C**) Complex III, (**D**) Complex IV, and (**E**) Complex V. (**F**) Heatmap of log₂ fold-change for individual proteins across OXPHOS complexes. Data are presented as mean ± SD. *n* = 5 mice per group. * *P* < 0.05 vs. indicated group.

### The activity of Complex IV is impaired following OVX

To assess whether changes in mitochondrial protein abundance translated into functional effects, we measured respiration in freshly isolated brain mitochondria. Glutamate-supported basal, State 3, and succinate-driven respiration showed no differences across groups (**Figure 6A–C**). Similarly, the oxygen consumption through specific Complexes I and II did not differ between groups (**Figure 6D and E**). In contrast, OVX significantly reduced the specific oxygen flux of Complex IV compared to the sham group (**Figure 6F**; *P*<0.05), a reduction that E2 treatment mitigated.

**Figure 6:**
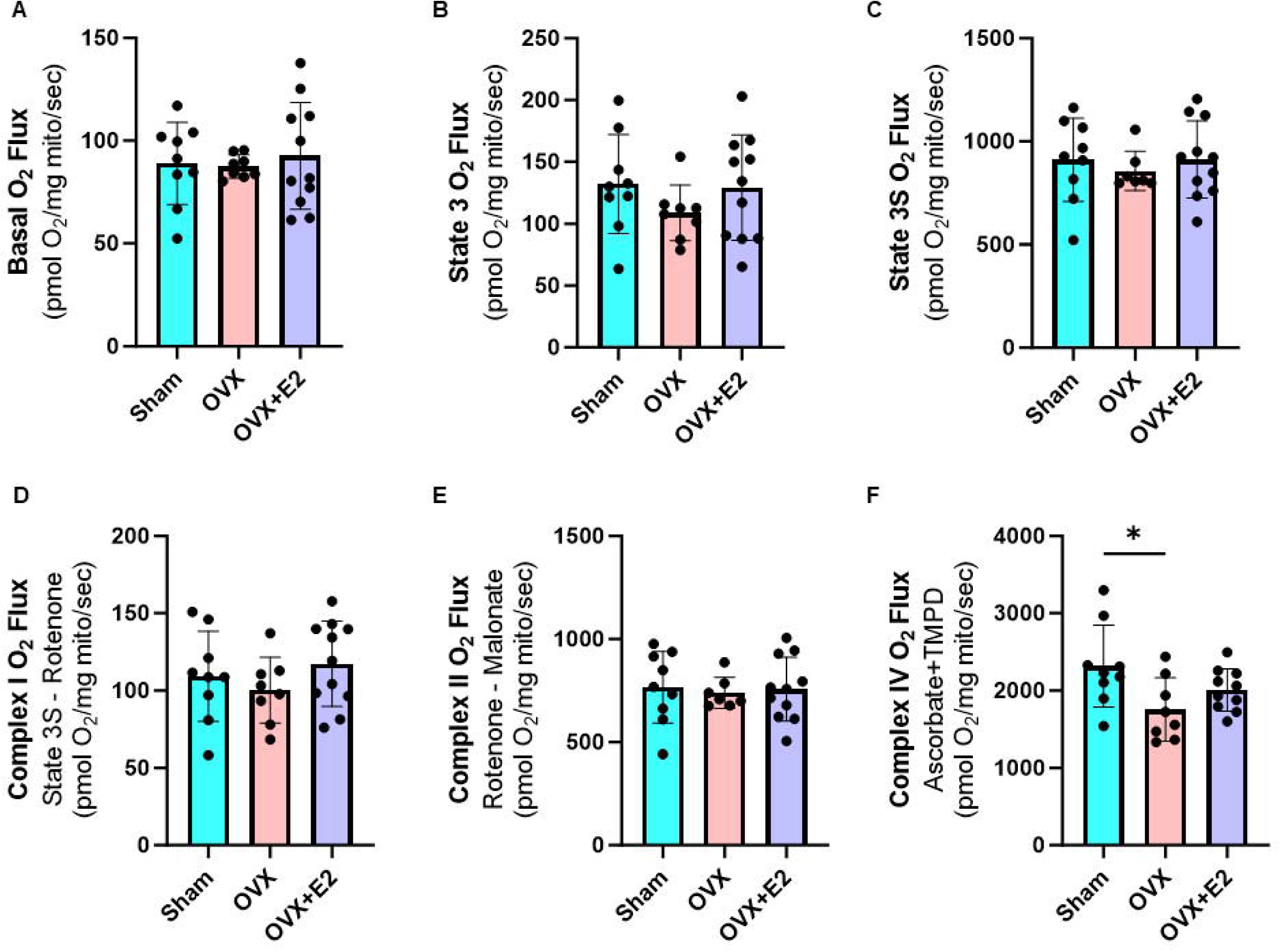
Isolated mitochondrial respiration and enzymatic activity: Glutamate-supported respiration measured in (**A**) basal, (**B**) State 3 (ADP-stimulated), and (**C**) State 3 Succinate (CI+CII). Enzymatic activity for (**D**) Complex I (inhibited by rotenone), (**E**) Complex II (inhibited by malonate), and Complex IV (supported by Ascorbate + TMPD). Data are presented as mean ± SD. *n* = 9/11 mice per group. * *P* < 0.05 vs. indicated group.

## DISCUSSION

Menopause is associated with cognitive decline and increased AD risk in women, in part due to the loss of estrogen and estrogen signaling [1, 2]. Hormone therapy (estrogen or estrogen plus progesterone treatment) has been reported to preserve cognition and mitigate AD pathology in women, but the molecular mechanisms remain unclear [5–9]. Prior work has demonstrated that E2 treatment following OVX enhances neuroplasticity, maintains cholinergic signaling, and supports mitochondrial respiration [12, 14–17]. Early proteomic studies using 2D-DIGE approaches demonstrated that acute E2 exposure increased proteins associated with synaptic signaling and mitochondrial function [16, 18]. Building on this, our study provides a deeper systems-level analysis of the impact of both OVX and OVX + chronic E2 treatment in the whole brain proteome of mice using data-independent acquisition (DIA) proteomics. We found that OVX decreased proteins involved in synaptic plasticity and brain metabolism, whereas E2 treatment restored synaptogenesis signaling by increasing proteins linked to presynaptic vesicle trafficking and cellular signaling. E2 also enhanced mitochondrial-associated proteins, rescuing BCAA and ketone metabolism pathways and restoring TCA cycle and OXPHOS proteins reduced by OVX. Importantly, these proteomic changes were accompanied by functional improvements, as E2 mitigated the OVX-induced decline in Complex IV protein abundance and enzymatic activity. Collectively, these findings identify E2 as a key regulator of synaptic plasticity and brain metabolic homeostasis.

Estrogen has been strongly associated with synaptic plasticity, as evidenced by previous reports that OVX reduces dendritic spine density while E2 treatment preserves it [29]. Consistent with this, prior studies have shown that E2 increases presynaptic markers, such as syntaxin, and postsynaptic markers, including spinophilin and PSD95, indicating enhanced synapse maturation [30]. At the proteomic level, acute E2 administration (24 hours) has been reported to increase vesicle proteins such as Stxbp1 and NSF [18]. Extending these findings, our results demonstrate that chronic E2 treatment following OVX restores presynaptic vesicle proteins, including Rab3a, Stxbp1, Snap25, and Rab5c, which were all reduced following OVX. These results suggest that E2 plays a role in maintaining the molecular mechanisms required for vesicle trafficking and neurotransmitter release. Interestingly, these changes align with proteomic studies in human AD brains, where mild-cognitively impaired (MCI) and AD patients show downregulation of vesicle trafficking proteins, including Stxbp1, Snap25, Stx1a, and Ap2a1 [31]. OVX in our model produces similar synaptic deficits, underscoring that E2 loss during menopause may recapitulate early AD-like synaptic changes. Among these proteins, Stxbp1 is of particular interest. Beyond being reduced in both OVX animals and AD patient brains, its knockdown in mice leads to deficits in both short- and long-term memory [32]. Thus, E2-mediated restoration of Stxbp1 and other vesicle proteins may represent a key molecular mechanism underlying the cognitive benefits associated with hormone therapy.

The transition to menopause in women is accompanied by significant changes in brain metabolism, characterized by a decline in glucose utilization and an increase in ATP production, which eventually plateaus after prolonged estrogen deficiency [33, 34]. Similarly, a rat perimenopause model recapitulates the reduction in glucose metabolism observed in humans [35], with several glycolytic intermediates (e.g., fructose-6-phosphate, 3-phosphoglycerate, pyruvate) shown to be reduced in the brains of estrogen-deficient rats [36]. Clinical studies further support that hormone therapy mitigates the decline in brain glucose metabolism caused by menopause compared to those who do not receive hormone replacement [34, 37]. Mechanistically, 4 hour E2 treatment has been reported to increase key glycolytic enzymes—including hexokinase (HK), phosphofructokinase (PFK), and pyruvate kinase (PK) [38]. In contrast, the present study, which employed chronic E2 replacement, did not detect widespread changes in the abundance of glycolytic proteins. However, we observed a selective increase in HK in OVX mice, which was normalized by E2 treatment, potentially reflecting a compensatory response to maintain glycolytic flux. More importantly, OVX resulted in reduced levels of the rate-limiting enzyme PFKP, which was restored with E2 treatment, suggesting a direct role for estrogen in sustaining glycolytic capacity.

Downstream of glycolysis, we identified alterations in mitochondrial pyruvate transport. OVX mice displayed a trend toward reduced pyruvate transport and oxidation, as evidenced by changes in the pyruvate dehydrogenase complex (PDH), its kinase (PDK), and phosphatase (PDP) subunits. Chronic E2 treatment reversed this trend, significantly increasing PDH-related proteins and restoring the potential for pyruvate flux into the TCA cycle. Notably, OVX also increased PDK3 expression, which phosphorylates and inhibits PDH, thereby blocking pyruvate oxidation and limiting the entry of acetyl-CoA into mitochondria. Restoration of PDH activity by E2 is consistent with prior work, which shows that acute E2 treatment upregulates PDH protein content within 24 hours [16]. Further, E2 treatment preserves PDH activity, which is reduced following the loss of E2 [4]. Collectively, these findings suggest that hormone therapy supports brain glucose metabolism not only by regulating the abundance of glycolytic enzymes but also by maintaining the critical step of pyruvate oxidation in mitochondria.

The current study also identified changes in alternative metabolic pathways, as OVX reduced both BCAA and ketone metabolism, which were restored with E2 treatment. BCAAs not only support nitrogen homeostasis and neurotransmitter cycling but also serve as auxiliary fuel sources by generating acetyl-CoA, acetoacetate, or succinyl-CoA [39]. Animal studies show that the postmenopausal transition is associated with reductions in brain BCAAs, potentially limiting BCAA metabolism [36, 40]. However, this reduction is restored with E2 supplementation [40]. Tracer studies in human and mouse brain slices confirm that BCAAs contribute to the TCA cycle through conversion to acetyl-CoA and succinyl-CoA, although iPSC-derived neurons from AD patients exhibit downregulation of BCAA metabolism [39]. The impact of menopause and E2 treatment on BCAA utilization, therefore, warrants further investigation. Ketone metabolism represents another critical alternative fuel source, particularly when glucose metabolism is limited. In rodent models, loss of cyclic E2 increases fatty acid uptake gene expression and reduces glucose metabolism, consistent with a compensatory shift toward ketone utilization within the brain [41]. In line with this, our findings demonstrate that E2 treatment following OVX enhances ketone metabolism by increasing Bdh1, a key enzyme in ketone oxidation. Given that ketogenic diets improve health span and memory outcomes, these results suggest that E2 may regulate brain energetics, in part, by promoting ketone metabolism. However, the role of Bdh1 in sustaining brain health remains poorly understood and requires further study.

Glucose, BCAA, fatty acid, and ketone metabolism converge on the TCA cycle by generating acetyl-CoA or succinyl-CoA, which are oxidized to provide reducing equivalents for OXPHOS. In the current study, OVX reduced the abundance of multiple TCA cycle proteins, consistent with prior reports of decreased intermediates such as succinic acid, citric acid, fumaric acid, and malic acid following E2 loss [40]. E2 treatment following OVX restored TCA protein content, including increases in Idh3g and Sdhc, in line with previous findings of elevated Mdh2 and Aco2 [16], and reversal of OVX-induced metabolite deficits [40]. Importantly, the reduced expression of TCA cycle genes in AD brain tissue correlates with both disease progression and cognitive decline [28], suggesting that preserving TCA function by E2 may be a critical mechanism for sustaining mitochondrial metabolism and protecting brain health.

During the menopause transition, women exhibit a decline in platelet mitochondrial COX activity (Complex IV), which correlates with reduced brain COX activity and cerebral hypometabolism [2]. Consistently, preclinical OVX models demonstrate impaired mitochondrial bioenergetics, including reduced complex I-supported state 3 respiration and complex IV enzymatic activity [4, 42]. Estrogen treatment, both acute and chronic, has been shown to restore state 3 respiration and enhance complex IV enzymatic function [4, 16]. In agreement, our study found that OVX impaired respiration through complex IV, which was mitigated by E2 treatment. At the protein level, OVX reduced complex IV subunits, whereas E2 restored Cox20 and Cox15. Cox15, required for heme A biosynthesis, is critical for maintaining complex IV function, and its mutations cause severe neurodegeneration, such as Leigh syndrome [43]. Notably, heme A levels are reduced in AD patients [44], suggesting Cox15 may represent a mechanistic link between estrogen, mitochondrial integrity, and AD pathology. Similarly, Cox20 knockdown *in vitro* reduces complex IV activity and mitochondrial bioenergetics [45], further underscoring the relevance of E2 in maintaining this essential respiratory complex.

Previous work also indicates that OVX reduces state 3 respiration without altering respiration following oligomycin treatment, suggesting a complex V rate limitation [4, 16]. In line with this, we observed a reduction in total complex V protein content following OVX, which was reversed with E2 treatment by restoring multiple subunits, including mAtp6, Atp5f1a, Atp5f1b, and Atp5mj. Consistent with our findings, acute E2 (24 h) treatment has been shown to increase complex V subunits Atp5a1, Atp5b, and Atp5c1 and enhance complex V activity [16]. Importantly, reductions in complex V subunits are also observed early in the brains of AD patients and animal models, where they are linked to diminished neuronal OXPHOS efficiency [46, 47]. Our findings suggest that estrogen supports ATP synthase integrity, potentially buffering against bioenergetic failure during E2 loss and early AD pathology. However, further studies are needed to define the precise mechanistic role of ATP synthase regulation by E2, including whether restoring complex V function directly translates into improved neuronal resilience and cognition.

This study is not without limitations. First, proteomic analyses were performed on whole-brain homogenates rather than region-specific samples. Because AD pathology is regionally selective, targeting vulnerable regions such as the hippocampus or frontal cortex would provide greater resolution on how OVX and E2 treatment alter brain proteomics. Second, the brain is composed of diverse cell types, including neurons, astrocytes, oligodendrocytes, and microglia; thus, the proteomic changes observed here may reflect mixed cellular contributions rather than cell-type–specific alterations. Single-cell or cell-type– resolved proteomic approaches could help clarify these effects. Third, while the OVX model captures features of estrogen loss, it also inhibits other ovarian functions, and it does not fully replicate the complexity of human menopause or AD pathology. Incorporating AD-relevant mouse models would provide greater translational relevance and allow direct investigation of whether the observed proteomic shifts increase or mitigate disease progression. Finally, this study focused on protein abundance. Still, it did not assess post-translational modifications (e.g., phosphorylation, acetylation, ubiquitination), which are critical regulators of protein function and key components of mitochondrial quality control mechanisms.

In conclusion, this study provides an in-depth proteomic analysis that reveals molecular mechanisms by which E2 treatment mitigates menopause-associated decline in brain health. We demonstrate that the transition from pre- to post-menopause-like condition via OVX alters the brain proteome, resulting in reduced synaptic plasticity and impaired key metabolic pathways. E2 treatment after OVX restores these processes by enhancing synaptic vesicle and signaling proteins, as well as rescuing metabolic pathways, including BCAA and ketone metabolism, the TCA cycle, and the OXPHOS system. Importantly, several subunits and assembly factors within these pathways, such as Cox15 in Complex IV and ATP synthase subunits, emerge as promising therapeutic targets. Targeting these proteins may provide strategies for hormone therapy mimetics that preserve mitochondrial function and energy metabolism in the absence of E2. Future studies should define cell-type–specific and region-specific proteomic changes, and test whether targeted interventions can translate these mechanisms into effective therapies for women at risk for AD.

## DATA AVAILABILITY STATEMENT

All data are found within the manuscript.

## ACKNOWLEDGMENTS

We would like to acknowledge Dr. Samuel G. Mackintosh and Dr. Stephanie Byrum at The University of Arkansas Medical Campus Proteomics Core (NIH R24GM137786 and P20GM121293) for their contribution to the proteomics analysis.

## CRediT

**SFB:** Writing – Original Draft, Data Curation, Writing Review. **EF**: Conceptualization, Data Curation, Formal Analysis, Investigation, Writing – Original Draft, Data Curation, Writing Review. **ZB:** Data Curation, Writing Review. **FBB:** Data Curation, Writing Review. **JA:** Data Curation. **AL:** Formal Analysis, Writing-Review, and editing. **HW:** Formal Analysis, Supervision, Writing-Review and editing **JT:** Conceptualization, Funding Acquisition, Methodology, Supervision, Writing-Review and editing. **BAK:** Writing – Original Draft, Conceptualization, Formal Analysis, Funding Acquisition, Methodology, Supervision, Investigation, Data Curation, Project Administration.

---

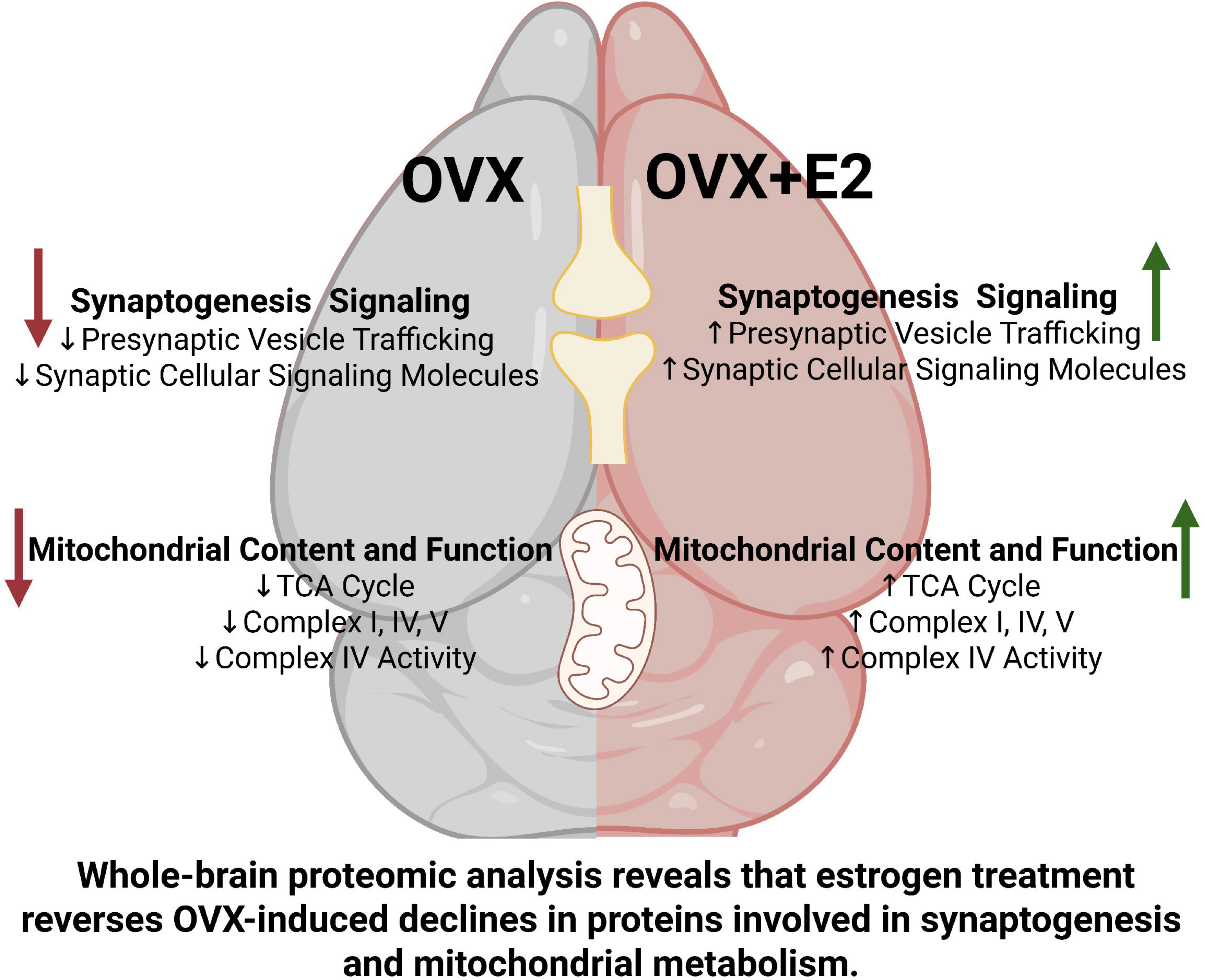

